# A summer in the Greater Paris: trophic status of peri-urban lakes shapes prokaryotic community structure and functional potential

**DOI:** 10.1101/2024.05.31.596803

**Authors:** P. Foucault, S. Halary, C. Duval, M. Goto, B. Marie, S. Hamlaoui, L. Jardillier, D. Lamy, E. Lance, E. Raimbault, F. Allouti, M. Trousselier, C. Bernard, J. Leloup, S. Duperron

**Author notes:** **Competing interest statement:** The authors declare that they have no known competing financial interests or personal relationships that could have appeared to influence the work reported in this paper. This work, as well as PF and MG grants were funded by the Agence Nationale de la Recherche project COM2LIFE (ANR-20-CE32-0006), MNHN and Sorbonne Université.

## Abstract

With more than 12 million inhabitants, the Greater Paris offers a “natural laboratory” to explore the effects of eutrophication on freshwater lakes within a relatively restricted area. Here, a four-month time-series was carried out during summertime to monitor planktonic microbial communities of nine lakes located within a ∼70 km radius around Paris (Île-de-France, France) of comparable morphologies, yet distinct trophic statuses (mesotrophic to hypereutrophic). The contribution of the trophic status and intra-summer variations were investigated on prokaryotic community structures (16S rRNA gene sequencing) and functions (shotgun metagenomics). Sampled lakes harbored highly distinct and diverse prokaryotic communities. Although their gene pool was quite stable and shared among lakes, taxonomical and functional changes were correlated. The trophic status appears as the main driver of both community structure and functional potential. When focusing on function involved in biogeochemical cycles, responses to phosphorus limitation (mostly polyphosphate-related processes) were highlighted between trophic statuses. Hypereutrophic lakes communities displayed the highest contrast and heterogeneity over time, suggesting a regime shift compared to lakes of lower trophic status. This study explores the influence of eutrophication on lakes’ microbiome ecology, by comparing for the first time the planktonic microbial communities and their functional potential of lakes of distinct trophic status in close vicinity over a summer season.

## Introduction

Over the last decades, lakes have been particularly affected by human activities, species invasion, increased surface temperatures and heat-waves associated with global change^1–4^. These add to the natural (*e.g.*, seasonal) variations and amplify eutrophication, ultimately leading to major changes in lake ecosystem functioning worldwide^5,6^. Increased eutrophication promotes blooms of phototrophs, that have tremendous consequences^7–9^ and are predicted to worsen over the next decades^2,7^. Understanding the link between eutrophication and lake functioning has thus become a priority for ecologists, environmental policy makers, as well as conservation scientists^5,10^.

Microbial communities are key contributors to ecosystem functioning^11^, quickly reacting to disturbances, and are thus often investigated to assess lake ‘health’. Eutrophication is a major driver of these communities^12–15^, promoting the growth of phytoplankton, including Cyanobacteria, as well as heterotrophic bacteria that degrade derived organic matter. However, the impact of eutrophication is often hard to disentangle from that of other variables (*e.g.*, lake morphology, land cover and uses^12,14^). Besides, communities variation with time (from days^16–18^ to years^19^) also needs to be accounted for. A few studies suggest that temporal variation of planktonic communities is affected by trophic status^20–23^. However, these time-series usually include a limited number of lakes, for example a single lake per trophic status. Moreover, higher trophic status reportedly leads to changes in community function^15,24,25^ (*e.g*, enhanced carbon and nitrogen fixation^24^), yet very few functional comparisons between trophic statuses are available.

To disentangle the link between eutrophication and lake functioning, we tested whether the trophic status drives the structure and the functional potential of prokaryotic communities during summer, when primary production is maximal. The Greater Paris (Île-de-France, France) offer a suitable playground. It is the 2^nd^ most populated European metropole (12 millions inhabitants over 814 km^2^) and harbors 248 artificial lakes according to Richardson *et al.*’s définition^26^), mostly old sand and gravel quarries, displaying distinct eutrophication levels^12,27,28^. It offers a “natural laboratory” to test the effects of distinct eutrophication levels on comparable lakes spread over a limited geographical area with limited effect of cofounding factors (*e.g.,* differences in climate, geological context, lake area and depth, pH). Here, nine shallow lakes of comparable morphologies displaying different trophic statuses were sampled monthly over the 2021 summer season. The structure and functional potential of prokaryotic communities were characterized by 16S rRNA amplicon and shotgun metagenome sequencing. We tested whether lakes different trophic status shapes the lakes microbial community structures and functional potential and altered their variability during summer.

## Material and Methods

### Sampling

Nine lakes were surveyed monthly from June to September 2021: Jablines (JAB), Vaires-sur Marne (VSM), Cergy large (CER-L), Cergy small (CER-S), Créteil (CRE), Bois-le-Roi (BLR), La Grande Paroisse (LGP), Champs-sur-Marne (CSM), Verneuil-sur-Seine (VSS). They are located within a ∼70 km radius around Paris (France; **Fig. 1A, S1**; see **Table S1** for coordinates), and were selected based on their similar area (7.3-91.0 ha), depth (3.5-10 m) and absence of stratification (**Table S2**). These are former sand and gravel quarries that were transformed into human leisure centers between the 1960s and the 1980s^12,27,28^.

**Fig. 1:**
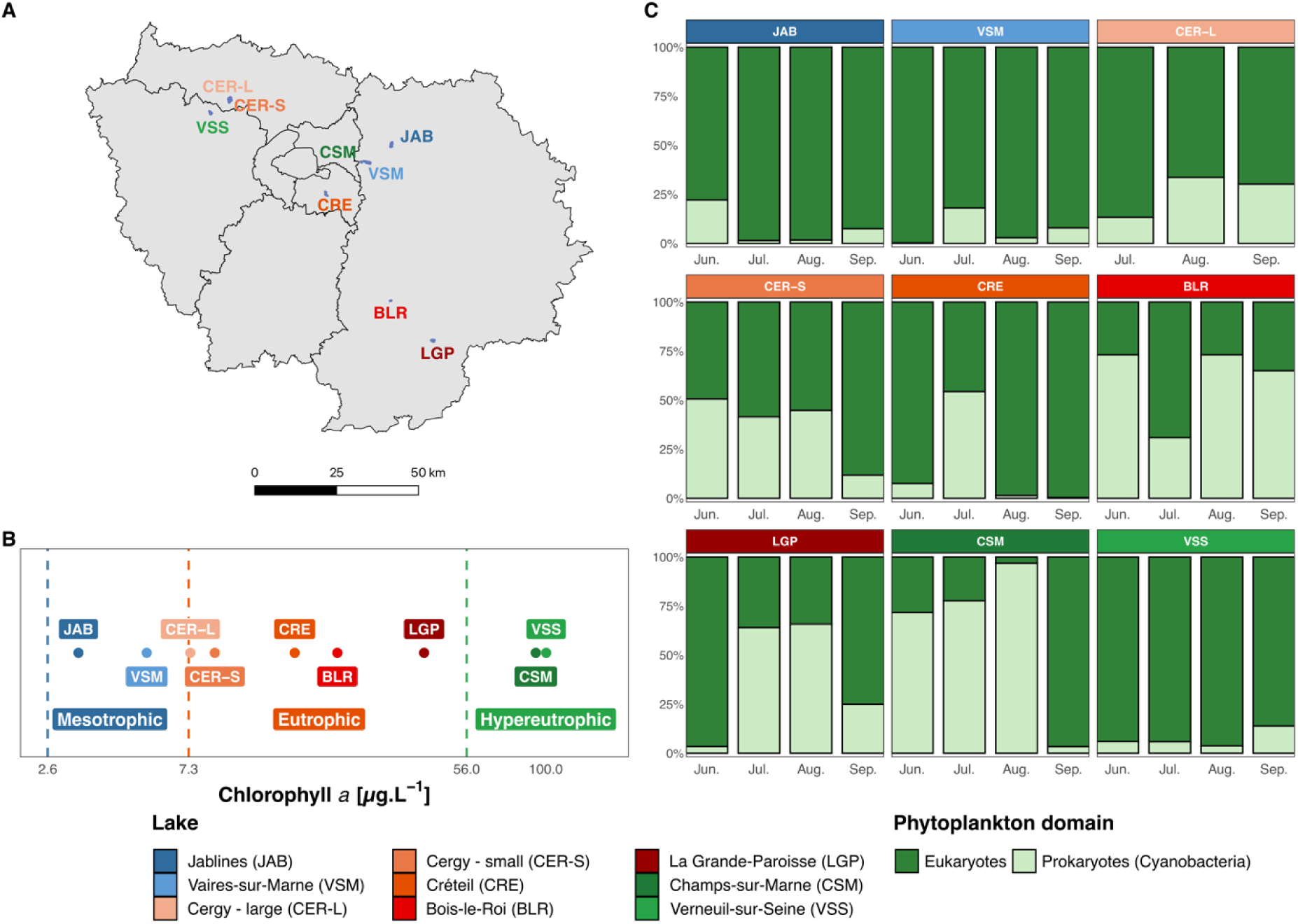
Lakes location, trophic status and phytoplankton community composition. **A:** Map of the Paris area (Île-de-France region), illustrating lakes location. **B**: Average Chl*a* concentration over the four months and trophic status based on Chl*a* concentration ranges proposed in the Carlson’s TSI guidelines (n=3 per lake for each month, except for VSM in July (n=2), 104 Chl*a* measures). **D:** Median of relative biovolume for each phytoplanktonic domain (104 samples).

In each lake, the water column was sampled at three mid-lake locations (labelled W1, W2 and W3, **Fig. S1**) to account for spatial heterogeneity. For each water column, 5 L were sampled using a Niskin bottle (WILDCO, USA) at 3 depths (∼0.5 m below surface, mid-depth and ∼0.5 m above the lake bottom), and then pooled together in equal volumes, forming a depth-integrated sample. A total of 105 samples were collected. CER-L could not be sampled in June. Samples were treated within one hour. Subsampling for Chlorophyll *a* (Chl*a*) concentration, phytoplankton composition, particulate carbon and nitrogen concentrations were obtained from unfiltered “raw-water”. For other subsamples, water columns were pre-filtered on 50-µm mesh to remove any large particles (*e.g.*, leaves and metazoan) prior to filtration and conditioning (referred as “pre-filtered water”). Conditioning and storage are described below.

### Physico-chemical parameters

Raw-water temperature and pH were measured on shore upon recovery (KS-2 MultiLine© probe, WTW, USA). Pre-filtered water was filtered onto 0.22-µm membranes (PES, Millipore Express, Germany). Eluates were collected in duplicate (2×12 mL) for nutrient analyses in polyethylene tubes, with a specific acidification (three droplets of 3% HNO_3_ solution) for orthophosphate (PO_4_^3-^ ions) analysis. Dissolved mineral nitrogen (NH_4_^+^, NO_3_^-^ and NO_2_^-^ ions) and PO_4_^3-^ concentrations were determined as described by Holmes *et al.*^29^. For particulate carbon and nitrogen concentration, around 1 L of raw-water (**Table S2**) was filtered onto 0.3-µm pre-combusted filters in duplicates (Sterlitec, USA). Filters and eluates were stored at - 20°C. Particulate carbon and nitrogen concentration were determined using a CHN Elemental Analyzer (NA1500 Series 2, Fisons, UK), values are expressed in µg and normalized by sampled volume.

### Chlorophyll *a*, phytoplankton identification and biovolume

The Chl*a* content (a proxy of phytoplankton biomass), was measured from 500 mL raw-water filtered onto 0.7-µm filters (GF/C, Whatman, UK), in triplicate, by spectrophotometry (Cary 60 UV-Vis, Agilent, USA), following Yéprémian *et al.*^30^. Phytoplankton composition was determined visually on lugol-fixed raw-water. Taxa identification and relative cell counts were performed under an inverted microscope (NIKON Eclipse TS100, Japan) based on the inspection of 200 to 400 random individuals per sample using the Utermöhl method^31^ (AFNOR 15204 standard). Cell biovolumes were based on previous reports from the Greater Paris lakes^12,32^. For taxa that were not in these reports, cell biovolumes were extracted from the 2017 HELCOM Phytoplankton Expert Group database^33^.

### Nucleic acids extraction

For each depth-integrated water column, 150 to 2,000 mL (**Table S1.1**) of pre-filtered water were filtered onto 0.22-µm membranes (PES, Millipore Express, Germany). Filters were flash-frozen in liquid nitrogen. Amplicon gene sequencing was performed on all 105 samples (W1 to 3, all lakes and dates except CER-L in June) and shotgun metagenomics was performed on 35 samples (W2 only, all lakes and dates except CER-L in June). Total DNA was extracted using the PowerLyzer PowerSoil DNA extraction kit (QIAGEN, Germany), including a prior bead-beating step (FastPrep-24 5G, MP Biomedical): five 30 s cycles (8 m.s^-1^) with 30 s pauses in-between (amplicon sequencing) and four 30s cycles with speed reduced to 6 m.s^-1^ (shotgun sequencing). Two extraction-blank controls were performed and incorporated into the 16S rDNA amplicon gene sequencing analyses.

### 16S rRNA gene amplicon sequencing

The V3-V4 region of the 16S rRNA-encoding gene was amplified using primers 341F (5’-CCTACGGGNGGCWGCAG -3’) and 806R (5’-GGACTACVSGGGTATCTAAT-3’, EMP Project^34^) using the following program: initial denaturation (94°C, 3 min); 35 cycles (94°C, 45 s; 55°C, 60 s; 72°C, 90 s); elongation step (72°C, 10 min). Products were sequenced on an Illumina MiSeq 250×2 bp platform (GenoToul, France). Amplicon sequence analysis was performed using the QIIME2 pipeline^35^ (version 2022.8). Amplicon Sequence Variants (ASVs) were obtained with the DADA2 algorithm: forward and reverse reads were trimmed at 230 and 225 bp, respectively, to keep a high phred quality score (median q > 30). The expected error rate was set at 2. Reads with a phred score < 20 and chimeras were discarded. ASVs were then affiliated taxonomically using the SILVA 138.2-99 SSU database^36^ and chloroplast- and eukaryote-affiliated reads were discarded. The analysis yielded 5,515 unique ASVs. Sample datasets were rarefied at 8,135 reads (lowest sample sequencing depth).

### Shotgun metagenomic sequencing

Genomic DNA from each W2 sample was sequenced (Illumina MiSeq 150×2 bp, GENOSCREEN, France) yielding 15.8 ±8 million paired-ends reads per sample. Sequence quality was checked (MetaWRAP pipeline^37^ (v1.3) and Multi-QC^38^ (v1.15)). Reads with a phred score below 20 were discarded. Human-associated reads were removed based on the GRCh38 human genome assembly. Samples were assembled individually using SPAdes^39^ (mode *meta*, v3.13.0) resulting in 1.8 ±0.8 million contigs per assembly (N50 = 383,761 ±217,731). Contig coverages were quantified in CPM units using Salmon^40^ (v0.13.1) in the *quant_bins* function of the MetaWRAP pipeline. The functional analysis was performed directly on the assembled contigs to investigate the gene-content at the community level, and avoid information loss associated with the MAGs binning process. Contigs were annotated taxonomically and functionally using CAT^41^ (v5.2) and Eggnog-Mapper^42^ (mode *prokaryota_broad*, v2.1.10). The final dataset consisted of a total of 7,994 annotated KEGG Orthologies (KOs).

A set of 28 marker genes were selected based on previous studies on aquatic prokaryotic communities^43–45^ and screened using Eggnog-Mapper annotations to further investigate processes related to carbon, nitrogen, phosphorus, sulfur and iron metabolisms. KO identifiers, corresponding processes, enzyme names and associated references are provided in **Table S3**.

### Statistical analyses

Statistical analyses were performed with R v4.1.3^46^ and RStudio. Mean Chl*a* concentration were computed by lakes and a Principal Component Analysis (PCA) was performed on other C-N-P nutrient parameters (scaled and centered TPC, TPN and NH_4_, NO_3_^-^ + NO_2_^-^, PO_4_^3-^ concentration values) using Vegan^47^ (v2.6-4). One sample (VSM, July, column W1) was discarded from all following analyses based on the aberrant measured Chl*a* concentration (**Table S2**). The correlation between the first PCA axis’ coordinates and the Chl*a* values was assessed by a Spearman correlation test (Rho coefficient (ρ), *cor.test*, Stats Rbase package v4.1.3). The taxa (ASV)- and gene (KO)-contents richness, evenness and Shannon indexes were computed using Phyloseq^48^ (v1.38.0) and Vegan. KOs and ASVs that were present within one given lake throughout all 4 sampled months were considered as part of its core gene- and taxa-contents, respectively. Month-to-month turnovers were computed for each lake (*turnover*, Codyn^49^ v2.0.5). In order to test whether taxa- and gene-content month-to-month turnovers differed among trophic status, while accounting for month comparison (month factor) and lake-specific effects (lake intercept), a linear mixed model (LMM) analysis was performed with the formula Y ∼ trophic status + month + (1 | lake), using the *lmer* function (Lme4^50^ v1.1-32 and LmerTest^51^ v.3-1.3).

To compare community dissimilarities based on gene- and taxa-contents, Principal Coordinate Analyses (PCoA) were performed using Bray-Curtis (BC) distances using Vegan. A Hellinger transformation was applied to the gene-content BC dissimilarity matrix to account for differences in metagenomic sequencing depth. The explanatory power of the ‘trophic status’, ‘month’ and their interaction term were tested using the *adonis2* (PERMANOVA) function of Vegan. Between-lake spatial distances were obtained using the *distHavsersine* function of Geosphere^52^ (v1.5-18). For each pairwise sample combination, the correlation between the gene- and taxa-contents BC dissimilarity values and between-lake distances was assessed by a Spearman correlation test (Rho coefficient (ρ), *cor.test*, package Stats Rbase v4.1.3). The intra-summer heterogeneity of each lake was visualized by plotting polygons representing the maximal area delimited by samples coordinates (in terms of gene- and taxa-content). Comparisons among lakes were performed on BC dissimilarity matrices (*betadisper*, Vegan). Differences between taxa- and gene-content intra-summer heterogeneity and BC dissimilarity value ranges were assessed by a LMM analysis (*lmer*, Lme4 and LmerTest packages) with the formula Y ∼ trophic status + (1 | lake).

SIMPER analyses were performed to identify ASVs or carbon, nitrogen, phosphorous, sulfur and iron biogeochemical cycles (BGCs) marker-genes that explained the differences between trophic status in their respective PCoAs (*simper*, package Stats Rbase v4.1.3). To avoid the detection of significant but rare ASVs, the analysis was performed on the subset of ASVs accounting for > 0.1 % of the overall dataset (148 ASVs) with the criterion of up to 70% of the cumulative explained dissimilarity (with an adjusted *p*-value < 0.001).

The correlation between the taxa- and gene-contents month-to-month pairwise dissimilarities (BC) within a lake was assessed by a Spearman correlation. For this comparison, only W2 water column samples were used because data was available for both gene- and taxa-contents.

All figures, except the Île-de-France and individual lakes maps, were created in RStudio using tidyverse^53^ (v2.2.0), ggConvexHull (v0.1.0), ggh4x^54^ (v0.2.3) and patchwork^55^ (v1.1.2). Legends were modified with Inkscape©. Values are displayed as “mean ± standard deviation” unless otherwise indicated.

### Sequencing data accession numbers

The 16S rRNA and shotgun metagenome raw reads were deposited into Sequence Read Archive (SRA, Project PRJNA1086840, see Table S1.1 and S1.2 for samples accession numbers). Scripts available at Github/pierrefoucault.

## Results

### Trophic status determination

The trophic status of the lakes was first described following Chl*a* concentration ranges from the Carlson’s trophic state index^56^ as oligotrophic (<2.6 *µ*g.L^-1^), mesotrophic (2.6 - 7.3 *µ*g.L^-1^), eutrophic (7.3 - 56 *µ*g.L^-1^) and hypereutrophic (>56 *µ*g.L^-1^; Fig. 1A). The complete index also uses Secchi disk depth, TP and TN that were not used here. Using the average summer Chl*a* concentration (Fig. 1B, Table S2), two lakes were classified as mesotrophic (JAB and VSM, respectively 3.3 ±2.4 and 5.4 ±4.2 *µ*g.L^-1^ Chl*a*), five as eutrophic (CER-L, CER-S, CRE, BLR, and LGP, respectively 7.39 ±2.14, 8.6±3.5, 15.9 ±7.8, 21.7 ±6.1 and 41.0 ±16.5 *µ*g.L^-1^), and two as hypereutrophic (CSM and VSS, respectively 92.9 ±24.6 and 100.0 ±115 *µ*g.L^-1^; Table S2). Photosynthetic eukaryotes dominated the phytoplankton (33.6 ±20.3 to 95.6 ±5.28 % of the phytoplankton biovolume over the four months; Fig. 1C): in the two mesotrophic (JAB and VSM), one eutrophic (CER-L) and one hypereutrophic lake (VSS). This highlights that whether eukaryotes or prokaryotes are the main Chl*a* producers is not a function of the trophic status. This is exemplified by both hypereutrophic lakes, VSS being dominated by *Ceratium* (Miozoa, 77.8 ±25.1% over July and August; Fig. S2C) while CSM was dominated by Cyanobacteria during 3 out of 4 months (June to August: *Aphanizomenon*, 43.8 ±40.5%, and *Dolichospermum*, 21.7 ±15.8%; Fig. S2C). BLR (eutrophic) was the only lake dominated by Cyanobacteria (mostly *Cyanocatena*) throughout all four months (60.5 ±20.1%; Fig. S2C).

To confirm our lake classification, a multivariate analysis was performed on nutrient parameters (Table S2). The PCA visually discriminated lakes along its first axis (Fig. S2A), which was highly and significantly correlated to Chl*a* concentrations (SPEARMAN, *p*<0.01and ρ 0.74; Fig. S2B, Table S4). Overall, classification in three categories based on the Chl*a* concentration from Carlson was thus supported by the nutrient-based classification, and is used throughout the study.

### Prokaryotic core gene- and taxa-contents

The assembly of the 35 individual metagenomes yielded 0.9×10^6^ to 5.5×10^6^ contigs, and 7,994 unique annotated prokaryotic KOs, of which 52% (4,123) were present in all lakes and months. All lakes displayed similar gene-content richness (6,036 ±331 KO) and Shannon diversity (7.0 ±0.15; Table S5). The core gene-content of each individual lake (*i.e.*, the genes present throughout all four months within a given lake) consisted of 5,138 ±271 KO. Over the four months, these core KOs represented most of the KO richness (73.5 ±3.1%) and were overwhelmingly dominant (99.8 ±0.09% of KOs abundance for a given lake; Fig. 2A, Table S5). This stability was confirmed by the low month-to-month KO turnover (15.4 ±2.2%; Fig. 2B). Thus, the core gene-content of a lake was both dominant and stable throughout the summer, whatever the lake and its trophic status.

**Fig. 2:**
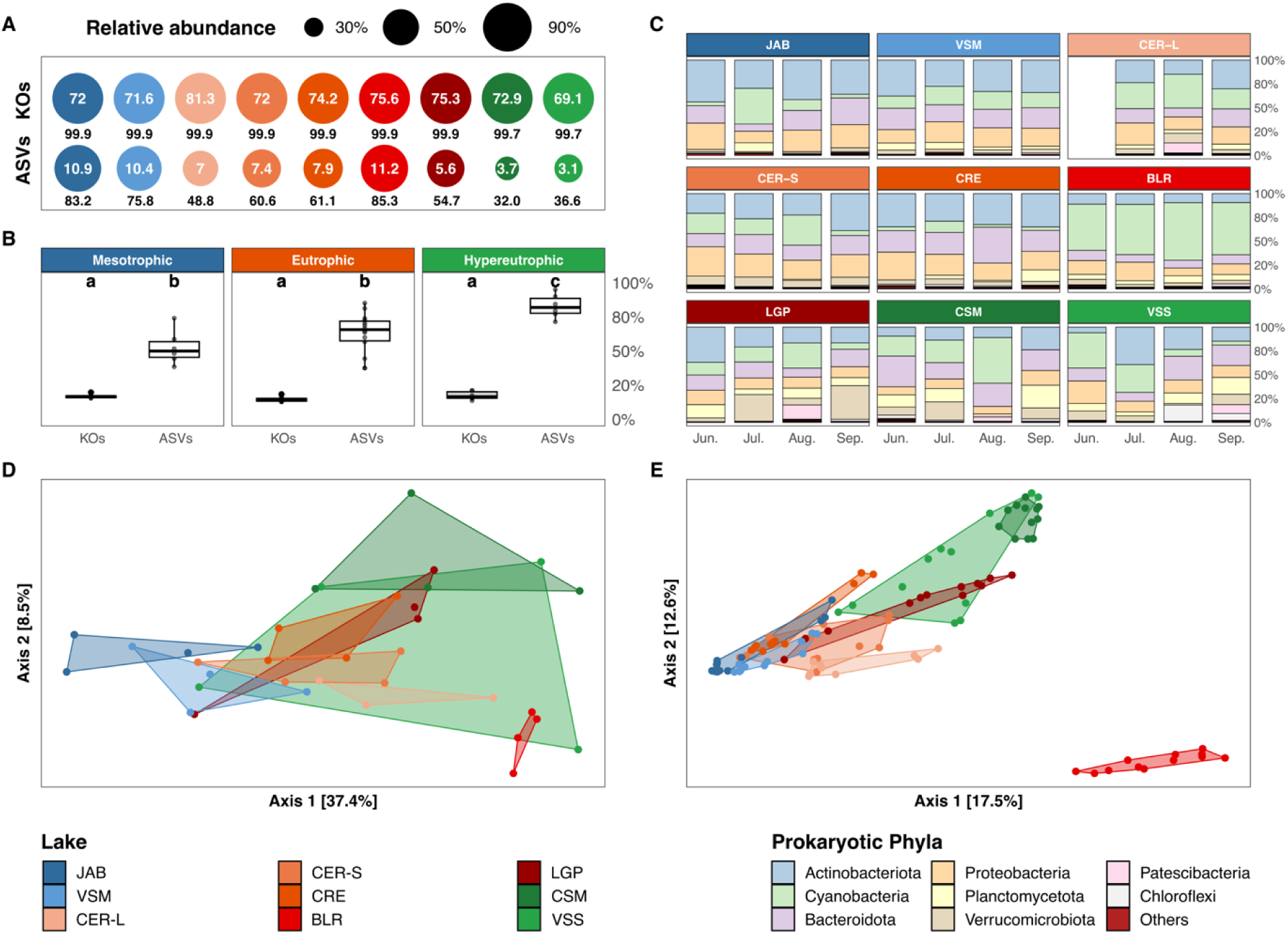
Prokaryotic gene- and taxa-contents composition (A-C) and structure (D-E). **A**: Individual lake’s core gene-(KOs) and taxa- (ASVs) contents. Circles area and the values below (in black) indicate the relative abundance of core KOs and ASVs reads versus all identified KOs and ASVs reads of a given lake. Inner circle values indicate the percentage of identified KOs and ASVs that belong to the core-content in a given lake. **B**: Month-to-month turnover (0 to 100%) of KOs and ASVs for each lake according to their trophic status. Letters indicate the significance of each trophic status (LMM). **C**: Taxa-content composition (phylum rank) as median proportion of total ASVs reads (104 samples). **D and E**: PCoA plots (BC dissimilarity), based on KOs (n=1 per lake for each month, 35 samples, **D**) and ASVs (**E**). Polygons represent the maximal area delimited by the samples coordinates of each sample for a lake. Lakes are colored according to their trophic status (see Fig. 1B).

Greater differentiation was observed for the taxa-contents. Indeed, only 5 out of 5,515 unique ASVs were shared among all prokaryotic communities (all water columns, 104 samples). Yet, the most abundant phyla were always Actinobacteriota (24.7 ±11.0%), Cyanobacteria (22.1 ±16.5%), Bacteroidota (18.2 ±6.91%), Proteobacteria (17.4 ±6.26%), Planctomycetota (8.0 ±5.25%), and Verrucomicrobiota (7.34 ±7.09%; Fig. 2C). The core taxa-content of each individual lake (*i.e.*, the ASVs that were present throughout all four months within the lake) accounted for a higher fraction of the ASVs richness in mesotrophic lakes (10.7 ±0.3%) compared to eutrophic (7.8 ±2.1%) and hypereutrophic lakes (3.4 ±0.4%; Fig. 2A, Table S6). The core taxa-content also accounted for a higher percentage of the ASVs relative abundance over the four months in mesotrophic lakes (79.5 ±5.2% *vs.* 62.1 ±13.8% and 34.3 ±3.2% for eutrophic and hypereutrophic lakes, respectively; Fig. 2A, Table S6). The month-to-month ASV turnover was also significantly lower in mesotrophic and eutrophic lakes compared to hypereutrophic lakes (respectively 53 ±13% *vs.* 63 ±15 and 83 ±9%; LMM, *p*<0.05; Fig. 2B, Table S7).

### Factors influencing the prokaryotic gene- and taxa-contents

Prokaryotic gene-contents of the lakes were segregated according to the trophic status along the first PCoA axis, with the exception of BLR (Fig. 2D). Lake’s trophic status had a significant effect (PERMANOVA, *p*<0.01 and R^2^ 0.21; Table S8), while no difference was observed according to neither sampling month nor spatial distance between lakes (Fig. S3A; Table S4, S7). The intra-summer heterogeneity (visually represented for each lake by a polygon linking the different sampling points in Fig. 2D) was significantly higher in hypereutrophic compared to eutrophic and mesotrophic lakes (LMM, *p*<0.05), the latter two categories displaying similar variabilities (LMM, *p*>0.05; Fig. S4A; Table S7). Heterogeneity was particularly high for VSS as illustrated by its large polygon area (Fig. 2D).

Overall similar trends were observed for the prokaryotic taxa-contents, with a segregation according to the trophic status (PCoA; Fig. 2E), which effect was significant (PERMANOVA, *p*<0.01 and R^2^ 0.17; Table S8). The sampling month as well as the interaction between sampling month and trophic status had significant, yet lower, contributions (PERMANOVA, *p*<0.01, R^2^ 0.07 and 0.13, respectively; Fig. 2E; Table S8). The intra-summer heterogeneity was significantly higher for hypereutrophic lakes (LMM, *p*<0.05) while similar between eutrophic and mesotrophic lakes (LMM, *p*>0.05; Fig. S4A; Table S7). The distance-decay relationship was significant but correlated poorly to the taxa-content dissimilarity (SPEARMAN, *p*<0.01 and ρ 0.22; Table S4). The taxa-contents from BLR appeared as a polygon offset and not overlapping with any other lake (Fig. 2E). Its taxa composition displayed both the lowest evenness (0.61 ±0.03) and Shannon diversity (3.3 ±0.2) over the summer period (Fig. S5; Table S6), and was dominated by a single cyanobacterial genus, *Cyanobium* (166 ASVs), which represented 55.1 ±5.65% of total reads throughout the four months *vs.* 13.5 ±11.3% in other lakes (Fig. S6). Similarly, on the gene-content dissimilarity plot, the BLR polygon also displayed very limited overlap (Fig. 2D).

A total of 34 ASVs significantly contributed to the difference between at least one of the pairwise trophic status comparisons (SIMPER analysis). The taxa-contents of mesotrophic and hypereutrophic lakes were set apart primarily by 7 Actinobacterial ASVs (*CL500-29* genus and *Hgcl* clade, together contributing to 25.1% of the difference) and 11 Cyanobacterial ASVs (*Cyanobium* and *Aphanizomenon* genera, 21.4% of the difference; Fig. S7A, D). Of lesser importance, seven Bacteroidotal and one Proteobacterial ASVs (ASV 3468) also contributed significantly to the difference between the mesotrophic and hypereutrophic lakes (respectively 12.9 and 10.6% of the difference for each phylum; Fig. S7A, D). The eutrophic status taxa-content was separated from the two other statuses by the lower abundances of aforementioned Cyanobacteria and Bacteroidota ASVs (Fig. S7B, C and D) and by ASVs displaying intermediate abundances between the mesotrophic and the hypereutrophic status (*e.g*., ASVs 3468 and 2304; Fig. S7D). Noteworthy, the higher abundance of four ASVs affiliated to Planctomycetota (either *Pirellula* or unassigned Pirellulaceae; Fig. S7D) and one Verrucomicrobiota ASV (*LD29*; Fig. S7D) contributed significantly to the difference between hypereutrophic versus eutrophic taxa-contents comparison (37.9% and 3.8%; Fig. S7C).

For each lake, gene- and taxa-contents month-to-month BC dissimilarities were compared (Fig. 3). Values were significantly correlated (SPEARMAN, *p*<0.01 and ρ 0.65; Table S4). The two hypereutrophic lakes CSM and VSS displayed both the highest gene- and taxa-contents dissimilarity values. (LMM, *p*<0.05; Table S7), compared to eutrophic and mesotrophic lakes, for which values were not significantly different (LMM, *p*>0.05; Fig. S4B, Table S7). Interestingly, BLR displayed the lowest range of both gene- and taxa-content dissimilarities (respectively 0.063-0.079 and 0.43-0.48; Fig. 3).

**Fig. 3:**
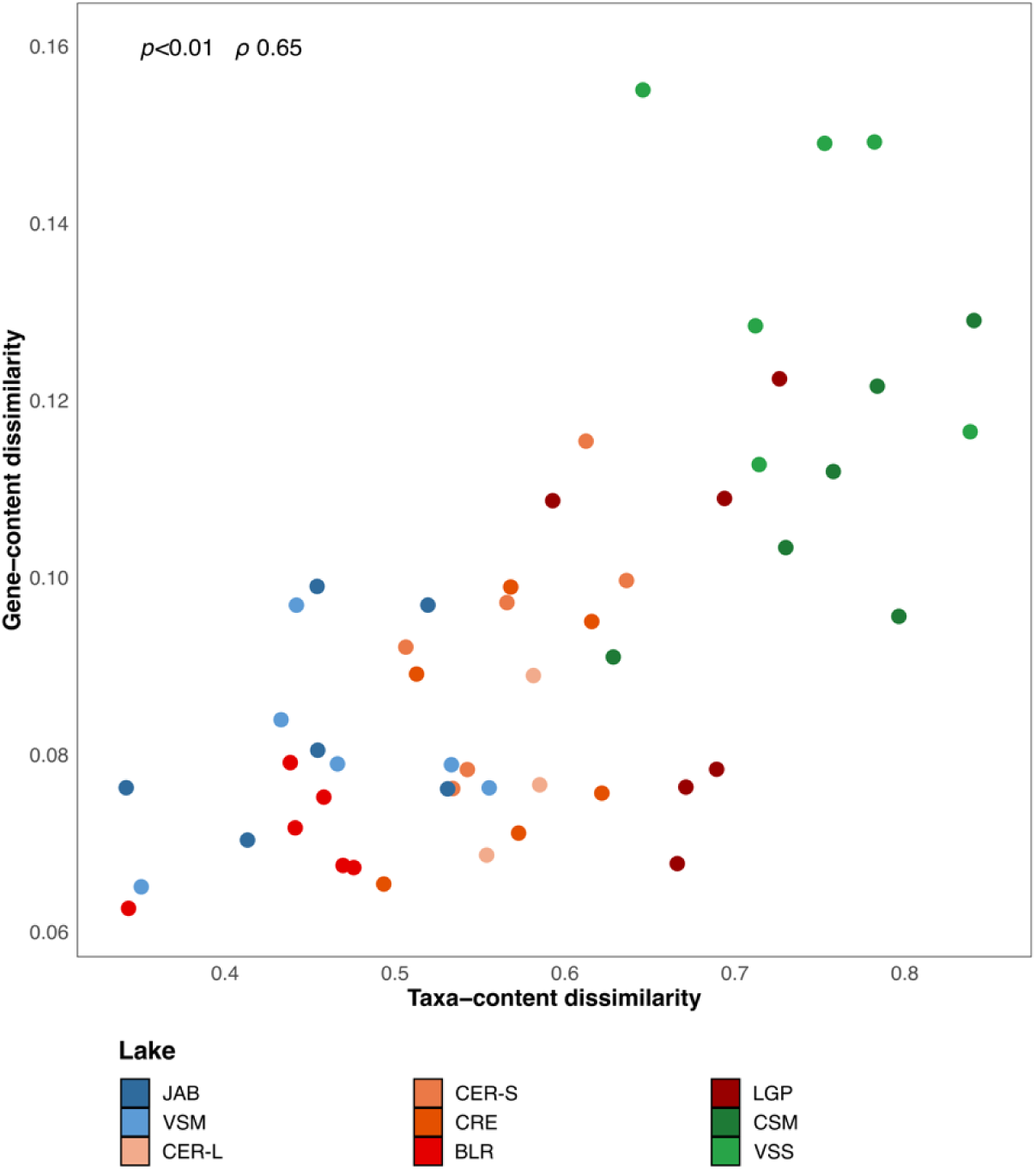
Gene-content *versus* taxa-content dissimilarities. Values on each axis correspond to month-to-month pairwise dissimilarities within a lake (y-axis: KOs, x-axis: ASVs, relationship assessed by Spearman correlation). Lakes are colored according to their trophic status (see Fig. 1B).

### Functional potential and BGCs marker genes contents

Relative abundances of the functional COG categories did neither show variation according to the different trophic status nor within a lake over the summer period (Fig. S8, Table S9). Almost 50% of contents grouped under 5 categories involved in metabolism and information processing: Translation, ribosomal structure and biogenesis (13.47 to 15.33%); Amino-acid transport and metabolism (10.42 to 10.66%); Energy production and conversion (9.97 to 10.33%); Replication, recombination and repair (8.84 to 8.88%) and Transcription (7.8 to 8.6%).

To explore the relationship between the trophic status and the functional potential related to the major biogeochemical cycles, 28 BGCs marker genes were selected (Table S3), together accounting for 0.7 ±0.3% of the total KO relative abundance. The abundances of these BGC marker genes clearly separated BLR on PCoA axis 1 (Fig. 4A). Moreover, lakes functional potentials were segregated according to the trophic status along PCoA axis 2 (PERMANOVA, *p*<0.01 and R^2^ 0.28; Table S8).

**Fig. 4:**
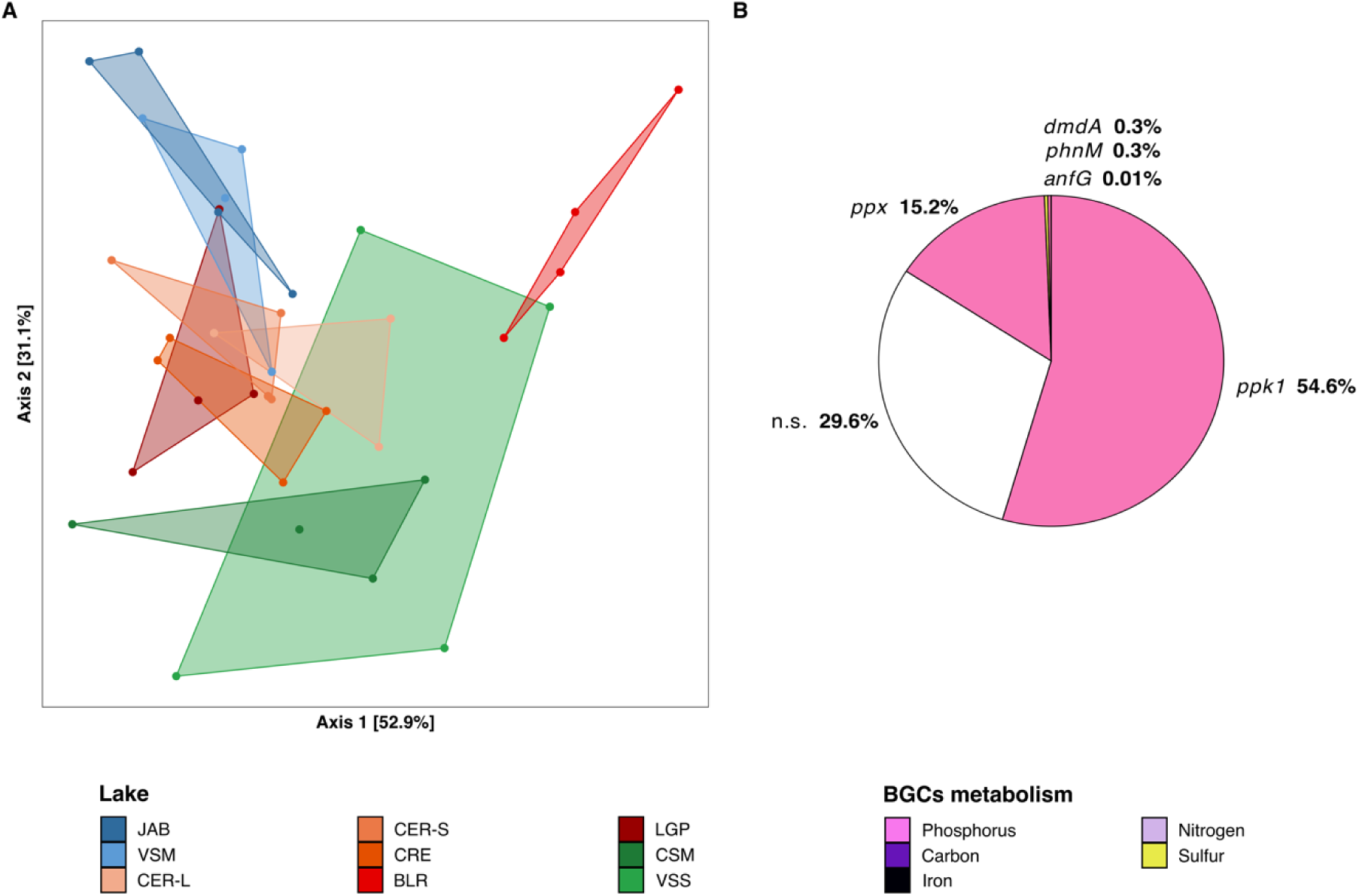
Prokaryotic gene-content structure based on a set of 28 BGCs marker genes. (carbon, nitrogen, phosphorus, sulfur and iron biogeochemical cycles)**. A**: PCoA plot (BC dissimilarity 35 samples). Polygons represent the maximal area delimited by samples coordinates for each lake. Lakes are colored according to their trophic status (see Fig. 1B). **B:** Percentage of the difference between mesotrophic and hypereutrophic communities, that is explained by BGCs marker genes (significant contribution (in %) of individual marker gene; **Table S10).**

Among the 28 BGCs marker genes, two genes involved in polyphosphate synthesis (*ppk1*) and hydrolyzation (*ppx*) were highlighted, both contributing highly and significantly to the differences, together explaining 69.8% of the mesotrophic - hypereutrophic status comparison (Fig. 4B, Table S10). Their relative abundances were higher in mesotrophic lakes (respectively 0.36 ±0.05 *vs.* 0.2 ±0.06% and 0.1 ±0.01 *vs.* 0.07 ±0.02%; Fig. S9, Table S3). Both genes were mainly detected on contigs affiliated to Actinobacteriota (72.4 ±13.1% and 62.3 ±13.9% of the *ppk1* and *ppx* KOs, respectively; Fig. S9). Noteworthy, the five other marker genes involved in phosphorus metabolism contributed significantly, yet to a lower extent (7.7% together) owing to their lower abundances, to the eutrophic *versus* hypereutrophic status comparison (Fig. S9, Table S3 and S10). The other marker genes contributed to a much lower extent (all less than 5% of observed difference; Tables S3 and S10). No difference in nitrogen metabolism marker genes was detected (Fig. S11, Table S10). Genes *psbA* (phototrophic activity), and *rbcL* (primary carbon fixation) involved in carbon metabolism were on average respectively six and fivefold more abundant in BLR compared to other lakes and mostly affiliated to Cyanobacteria (>90%; Fig. S10).

## Discussion

All nine lakes were located in close vicinity from one another around Paris with comparable features^12,27,28^, yet they were categorized into three distinct trophic status, providing an opportunity to test how trophic status affects prokaryotic community structure and functional potential during summer, with limited confounding factors compared to larger-scale studies.

### Peri-urban lakes display a stable gene-content throughout the summer despite taxa changes

The composition of the phototrophic communities varies among lakes and during the summer season. Most of the time, it is dominated by eukaryotes, BLR being the only lake where Cyanobacteria dominate throughout all four months. In the prokaryotic communities, four main bacterial phyla dominate, namely Actinobacteriota, Proteobacteria, Cyanobacteria and Bacteroidota, all being commonly reported in freshwater lakes^65–68^. At the ASV level though, the core taxa of a lake represent only a small fraction of its taxa richness, indicating that very few taxa are consistently found throughout the summer and emphasizing the high variability of taxonomic composition in a given lake. This is not unprecedented, as only 1.6% of prokaryotic taxa were shared by five ponds located within a 10 km radius in the southwest of Greater Paris^61^. Several previous studies have also highlighted the higher taxa turnover in freshwater ecosystems compared to soil for bacteria^62^ and protists^63^, and pointed to higher microbial predation pressure in lakes as a possible explanation^64^. On the other hand, the gene-content of each lake is stable in our study. Indeed, core genes present throughout all four months are overwhelmingly dominant in both richness and abundance, indicating much lower functional than taxonomic turnover. Besides, over half of all genes identified in this study occurred in all lakes and months, suggesting that despite large variations in taxa contents, a mostly shared set of functions underlays prokaryotic processes during the summer season. Greater variation of taxa-contents compared to gene-contents (tenfold) was recently observed between freshwater lakes spread over all Canada, sampled once during summer^15^. Altogether, our findings thus reflect trends previously observed at various spatial scales and confirm that studying microbial communities in spatially close lakes is a relevant approach to tackle microbial ecology questions if one wants to limit the influence of confounding climatic and geographic factors.

Our results also support the hypothesis of high functional redundancy among taxa in Greater Paris lakes, *i.e.* functions can be carried out by multiple taxa and are much more conserved than the taxa themselves^65,66^. This suggests that microbial diversity contributes to “buffering” the microbial ecosystem functioning, as an insurance against stressors^67^ (*e.g.*, contaminations, species invasion, raising surface temperature). However, changes in community structure and functional potential are still correlated, indicating that functional redundancy, if important, is not complete^44,68^. Strong links between changes in the community structure and their functional potential have been reported in freshwater lake ecosystems^15,24,69^ as well as in marine habitats^44^. This relationship was found even stronger when functionally-unannotated genes were included in such comparison^44^, indicating that much of the variation occurs in functions that are not yet properly characterized^70^. As we annotate the “common” functions, many of which are central cellular processes shared by all organisms, the high functional redundancy hypothesis must be taken with a grain of salt. Also, one next step would be to measure functions expression using metatranscriptomic approaches. Because gene expressions are highly sensitive to changes over a short time scale, typically hours^71^, they do not mirror differences highlighted from metagenomic analyses, and must rely on appropriate sampling frequency that is different from the monthly frequency employed herein.

### Trophic status is the main driver of prokaryotic community structure and functions

Only a handful of studies have investigated the relationship between freshwater lakes eutrophication and their planktonic prokaryotic communities by comparing various trophic statuses over time^20–23^. Here, the prokaryotic community structure correlates with the trophic status of nine lakes located in the Greater Paris, even when accounting for their intra-summer heterogeneity or the relatively short distances between lakes. Higher Chl*a* concentrations of hypereutrophic lakes (CSM and VSS) reflect the dominance of blooming taxa including cyanobacterial genera *Dolichospermum* and *Aphanizomenon*^7^ or the eukaryotic genus *Ceratium*^72,73^. These phototrophs provide an autochthonous source of organic matter to those lakes. This probably explains the higher abundances of heterotrophic bacterial phyla known for their ability to degrade phytoplankton-derived organic matter, including Verrucomicrobiota (*e.g.,* clade *LD29*) and Planctomycetota (*e.g.,* Pirellulaceae)^74–76^. Clade *LD29* is for example abundant in the highly eutrophicated Baltic Sea and in mesotrophic to eutrophic lakes^77^, where it lives within the phycosphere and degrades polymers^78^, while taxa belonging to the Pirellulaceae have been shown to degrade sulfated polysaccharides derived from cyanobacterial mucilage^79–81^. In contrast, mesotrophic lakes (JAB and VSM) communities were characterized by ASVs belonging to freshwater Pelagibacteraceae (Alphaproteobacteria Clade III), able to thrive in environments with low phytoplankton biomass and low nutrient availability^82,83^, and Actinobacteriota, able to degrade allochthonous organic matter, notably complex plant- (*e.g.*, lignin, cellulose, xylan)^84–86^ and zooplankton-derived polymers (*e.g.*, chitin degradation by-products)^87^. Those findings are overall congruent with various studies in terms of which groups vary according to the different trophic status^20–24,88,89^. However, previous studies either monitored prokaryotic communities of freshwater lakes in one-shot sampling campaign during summer^24,88,89^ (peak of primary production), or focused on inter-seasonal variations^20–23^, omitting intra-seasonal variability (*e.g.*, one campaign in April and August^21^). Furthermore, selected lakes usually displayed more distinct morphometric properties than here (*e.g.*, lakes depths from 12 to 58 m^20^), or featured only two trophic statuses (*e.g.*, mesotrophic *vs.* eutrophic lakes^21^). For example, Aguilar *et al.*^22^ monitored the prokaryotic communities of one oligotrophic and one mesotrophic alpine freshwater lake monthly for over a year and reported that changes in prokaryotic community structures between these two lakes were comparable in summer, and contrasted in winter. This very interesting study is however difficult to compare with ours owing to the very different alpine context (*i.e.*, the altitude higher than 900 m, the ice-covered period, lack of surrounding human activities) and lakes lower trophic status.

Besides its influence on taxa, trophic status also influenced, to a lesser extent though, the overall functional potential encoded by metagenomes. When considering functions involved in BGCs, trophic status mostly influenced processes related to phosphorus metabolism, the typically limiting nutrient in freshwater lakes^90–92^. In our study, mesotrophic lakes were characterized by higher abundances of genes involved in polyphosphate metabolism harbored by Actinobacteria (*ppk1* and *ppx* genes). Polyphosphate formation is associated with phosphorus limitation in marine ecosystems^93,94^ and, while also well-documented in Cyanobacteria^95–97^, is well described in genomes of Actinobacteria^86,98^. Hypereutrophic lakes (particularly CSM) were characterized by higher abundance of phosphonate utilization genes (*phnM* and *phnD* genes) affiliated to Proteobacteria and Cyanobacteria. Phosphonate is an organic source of phosphorus found as a xenobiotic in polluted aquatic ecosystems^99–102^ or derived from organic matter degradation. Heterotrophic bacteria able to degrade phosphonate compounds were recently shown to be abundant in the phycosphere of bloom-forming Cyanobacteria^103^. Altogether, and despite overall high functional redundancy, functions associated with phosphorus metabolism might be among those that are affected by trophic status. Other seasons have been reportedly associated to higher prokaryotic carbon, nitrogen, sulfur metabolism variability in marine and coastal ecosystems^44,45^, indicating that these should be explored also during other seasons in lakes from the greater Paris.

### The BLR lake illustrates atypical stable dominance of Cyanobacteria

The microbial community of lake BLR appears as an outlier in our study. The phytoplanktonic community is dominated throughout the summer by *Cyanocatena*^104,105^, a genus of small Cyanobacteria described in other artificial lakes, for example in an old gravel pit lake near Bratislava (Slovakia)^106^ or in Lake La Preciosa^107^ (Mexico). The sub-50 µm prokaryotic community was also dominated by small Cyanobacteria, namely *Cyanobium*^104,108^, enhancing the potential for photoautotrophy (*rbcL* and *psbA* genes) and nitrogen cycle related processes (*ureC* and *narB* genes), although not its fixation. Stable dominance of a limited diversity of Cyanobacteria (*Cyanocatena* and *Cyanobium*) could explain the overall stability observed in prokaryotic taxa and functions^17^. None of the environmental parameters analyzed herein explains this greater stability compared to other lakes. A previous study found that the surface picophytoplanktonic community of Lake Erie was dominated by strains closely related to freshwater *Cyanobium*^109^. Authors suggested the higher total dissolved phosphorus and lower silicate concentrations of Lake Erie, compared to other Great Laurentian Lakes, as possible explanations, but in our case, BLR does not differentiate from other lakes in terms of PO_4_^3-^ concentration. The stable dominance of *Cyanocatena* and *Cyanobium* in their respective size-fractions, and their lower abundances in the eight other lakes sampled in this work, suggest that unidentified controlling factors might be at play. Additional biotic factors (*e.g.*, microbial eukaryotes, zooplankton) should be investigated. Indeed, the phytoplankton and zooplankton diversity have been shown to be positively correlated in the Laurentian Great lakes^110^, and higher zooplankton richness has been linked to greater phytoplanktonic community stability in mesocosm experiments^111^. Increasing cyanobacterial abundance in freshwater ecosystems around Cracow (Poland) during summer has been negatively correlated to the functional richness of the zooplankton community^112^, supporting a possible link. Whatsoever, the stability of the BLR community is a great opportunity to study the interactions between multiple trophic levels and the microbial loop and their consequences on community functioning.

### Does the hypereutrophic status induce a regime shift for prokaryotic communities?

The two hypereutrophic lakes CSM and VSS exhibit the lowest number of core taxa, the highest taxa turnover, and intra-summer variability in taxa- and gene-contents. These features set them apart from mesotrophic and eutrophic lakes. These two lakes also display differences in community structure and functional potential between each other. The dominance of distinct phytoplanktonic domains could be one explanation for these differences, with Cyanobacteria dominating in CSM (3 out of 4 months) while eukaryotes dominate in VSS. Indeed, different phytoplanktonic taxa release different quantity and quality of organic matter and nutrients in freshwater and other aquatic ecosystems, leading to distinct bacterial communities^113–115^.

High community heterogeneity in hypereutrophic lakes is going against the common expectation of increased community homogeneity in nutrient-rich ecosystems, as documented for example for phytoplankton^32^ or Cyanobacteria and micro-eukaryotes in lakes during the last decades^116,117^. On the other hand, our results are in agreement with other works showing increased heterogeneity with higher trophic status when comparing freshwater lakes planktonic microbial communities across space^88,118^ and time^20,21,23^. There is recent evidence that protist communities’ heterogeneity also increases with freshwater lakes trophic status from a large scale study in Canada during summer^119^. Some studies point to the potential role of viruses, which may display distinct strategies depending on the trophic status (notably the shift between lysogeny and lytic cycle), although results are not clear-cut^120–122^. Thus, we hypothesize the existence of an alternative regime associated with hyper-eutrophication for microbial communities. In this regime, prokaryotic taxa and functions would display higher variability compared to those from lower trophic statuses. Whether this prokaryotic community “regime shift” occurs only during the summer as well as the underlying causes need to be further explored.

Analyzing overall comparable lakes spread over a limited area near Paris, yet with trophic status ranging from meso-to hypereutrophic, allowed us to show that trophic status has an impact on community structure and functional potential in summer. Functional potential is much more stable than taxa composition within each lake, most of it being shared among all lakes. High eutrophication levels are sometimes assumed to be irreversible, so whether their driving effect and the hypothetical “regime shift” in hypereutrophic lakes continue in periods of lower primary production, such as winter and spring, needs to be tested. Besides, the identification of one eutrophic lake displaying very stable communities in comparison to other suggests that trophic status alone cannot explain all observations. For this, peri-urban areas such as the Greater Paris, in which lakes of various trophic status occur in close vicinity, provide excellent settings.

## Abbreviations

Chl*a*: Chlorophyll *a*
ASV: Amplicon Sequence Variant
KO: KEGG Orthology
BC: Bray-Curtis
BGC: Biogeochemical Cycle
PCA: Principal Component Analysis
PCoA: Principal Coordinate Analysis

## Acknowledgements

We thank the COM2LIFE consortia for their participation on the fieldwork (authors of this work, Siham MESLI, Maëlle JARNO, Alison GALLET) and especially Haeitz ALOUI for the administrative work. We thank the leisure’s centers directors for their monthly access approval. We thank the PCIA plateform for their computational facility (Plateforme de Calcul Intensif et Algorithmique, MNHN, CNRS, UAR 2700 2AD, Paris, France), the GENOTOUL and GENOSCREEN sequencing facility. We thank the PARI platform (Plateforme d’Analyse Haute Résolution, IPGP, UMR 7154, CNRS, Paris, France) and the MIO-PACEM platform (P. Raimbault, Plateforme Analytique de Chimie des Environnements Marins, Aix Marseille Université, CNRS, Marseille, France) for nutrient mesurements.

## Data availability

All 16S rRNA gene amplicon sequencing and shotgun metagenomic raw reads were deposited into the Sequence Read Archive (SRA) database under the Project PRJNA1086840 (see Table S1.1 and S1.2 for individual sample SRA accession numbers). Scripts available at Github/pierrefoucault.

## Authorship contribution statement

PF: Investigation, Data curation, Formal analysis, Writing – original draft. SH: Conceptualization, Investigation, Writing - Review & Editing, Supervision, Funding acquisition. CD, MG: Investigation. BM, LJ, SH: Conceptualization, Investigation, Writing - Review & Editing, Funding acquisition. DL, EL, ER: Investigation. FA: Resources. MT, CB: Conceptualization, Writing - Review & Editing, Funding acquisition. JL, SD: Conceptualization, Investigation, Writing - Review & Editing, Supervision, Project administration, Funding acquisition.

