## Supplementary figures for "A summer in the Greater Paris: trophic status of peri-urban lakes shapes prokaryotic community structure and functional potential"

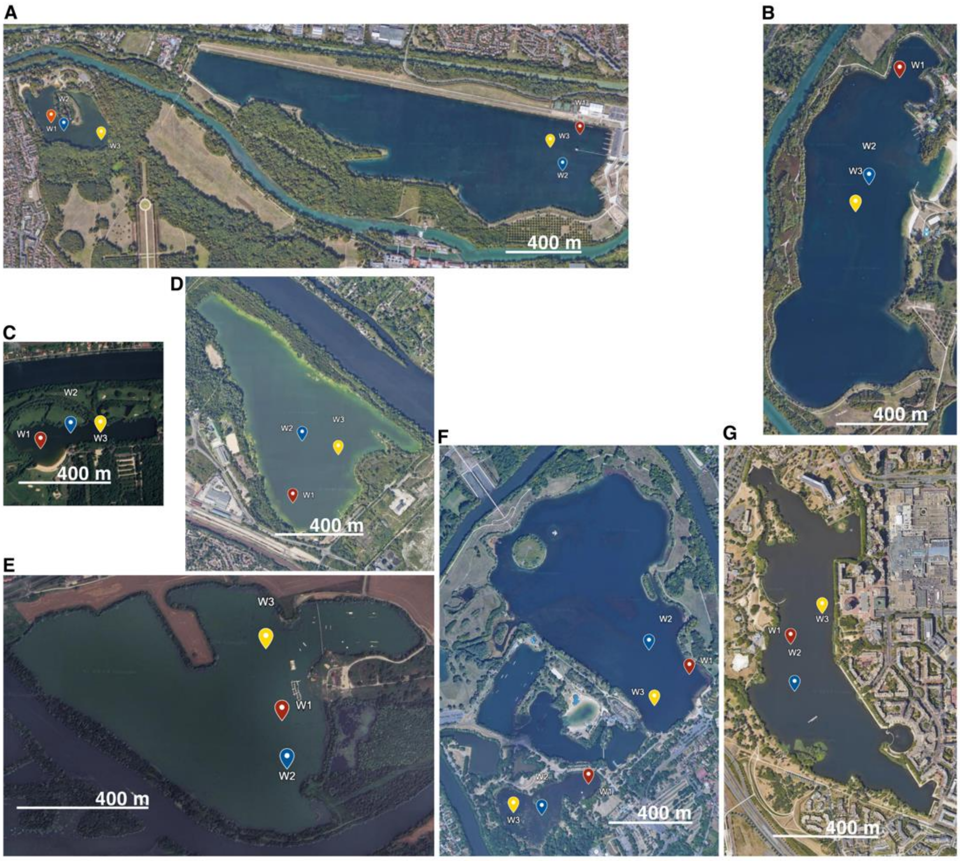


**Fig. S1: Satellite view of the lakes.** Images were taken from Google Earth © displaying the three sampling coordinates within each lake (see **Table S1.1** and **S1.2** for coordinates) : CSM (**A**, on the left; 3.6 m depth and 9.7 ha), VSM (**A**, on the right; 4.7 m depth and 86.7 ha), JAB (**B**; 7.4 m depth and 76.9 ha), BLR (**C**; 2.6 m depth and 7.3 ha), VSS (**D**; 4.9 m depth and 45.8 ha), GDP (**E**; 3.75 m depth and 51.9 ha), CER-S (lower **F**; 2.5 m depth and 10.5 ha), CER-L (upper **F**; 5.75 m depth and 91.0 ha) and CRE (**G**; 5.1 m depth and 40.1 ha).

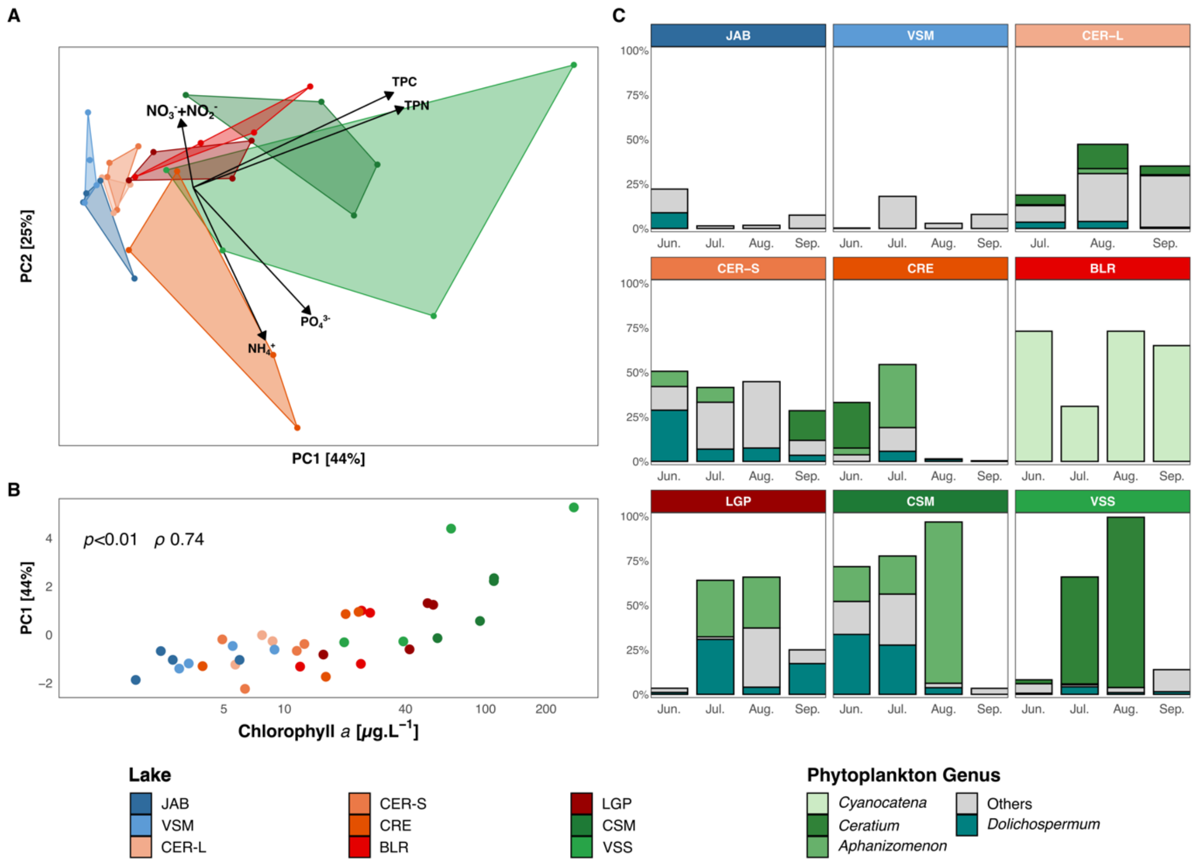


**Fig. S2: Supplementary analysis of abiotic and biotic parameters. A:** PCA plot based on nutrients parameters (TPC, TPN, $PO_4^{3-}$, $NH_4^+$, $NO_3^-$+$NO_2^-$, Table S2). Polygons represent the maximal area delimited by samples coordinates for each lake. **B:** Relationship between the Chl*a* concentration and the PC1 coordinates, assessed with a Spearman correlation. **C**: Median relative biovolume of *Ceratium* (Miozoa) and all Cyanobacterial genera (grouped in "Others" if not affiliated to *Cyanocatena*, *Dolichospermum* or *Aphanizomenon*, 104 samples). Lakes are colored according to their trophic status (see **Fig. 1B**).

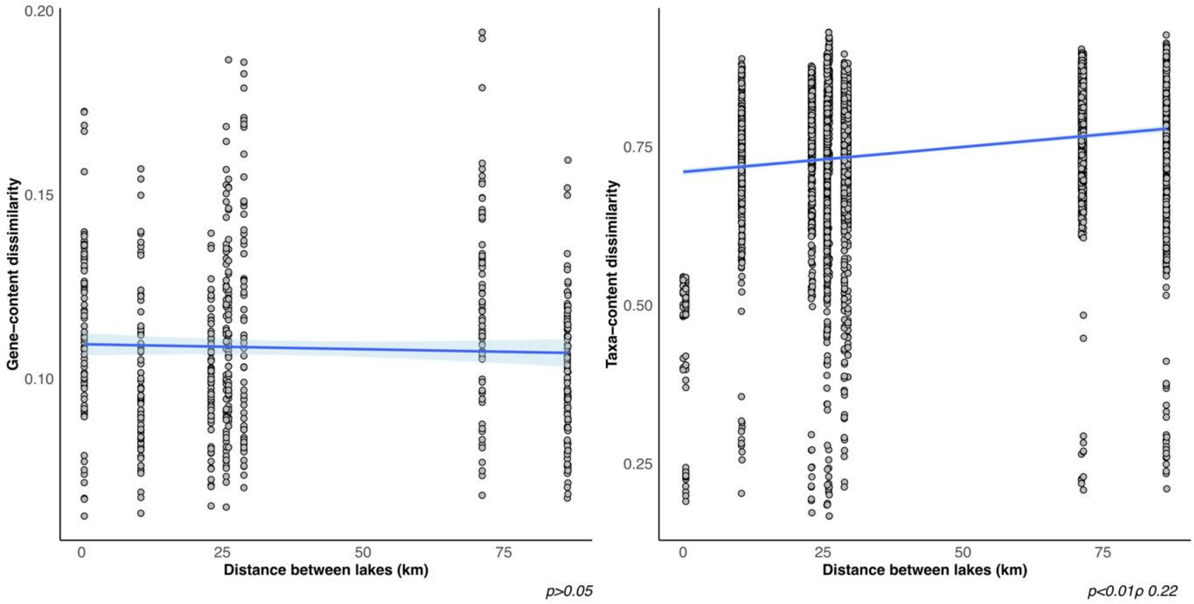


**Fig. S3: Influence of the spatial distance between lakes on the gene- and taxa- content dissimilarities. A and B:** Relationship between the gene- (KOs, 35 samples, **A**) and taxa-content (ASVs, 104 samples, **B**) dissimilarities (BC) and the distances between lakes (in km). Relationship significances are assessed by Spearman correlation.

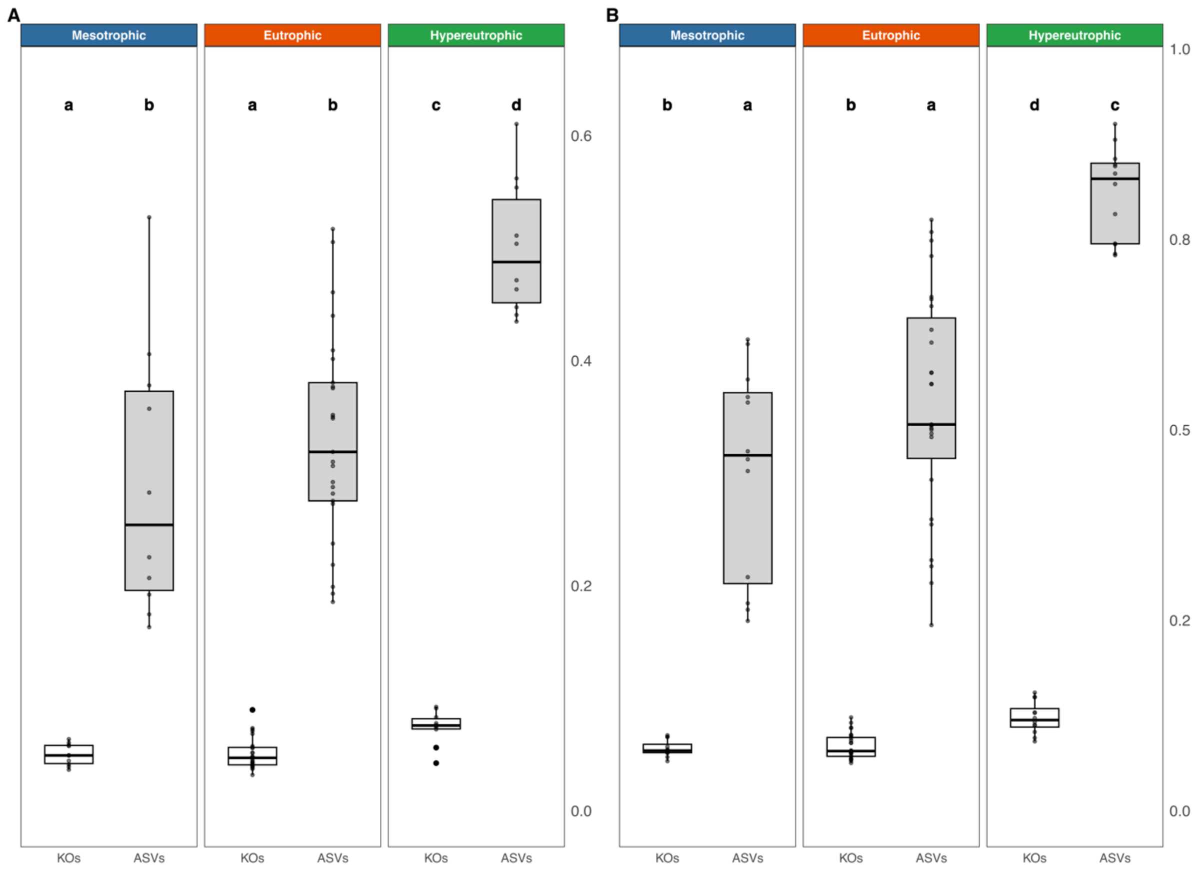


**Fig. S4:** Intra-seasonal heterogeneity (**A**) and overall dissimilarity value ranges (**B**) based on gene-content (KOs, white boxplots) and taxa-content (ASVs, grey boxplots) BC dissimilarity for each lake according to their trophic status. Letters indicate the significance of each trophic status (LMM).

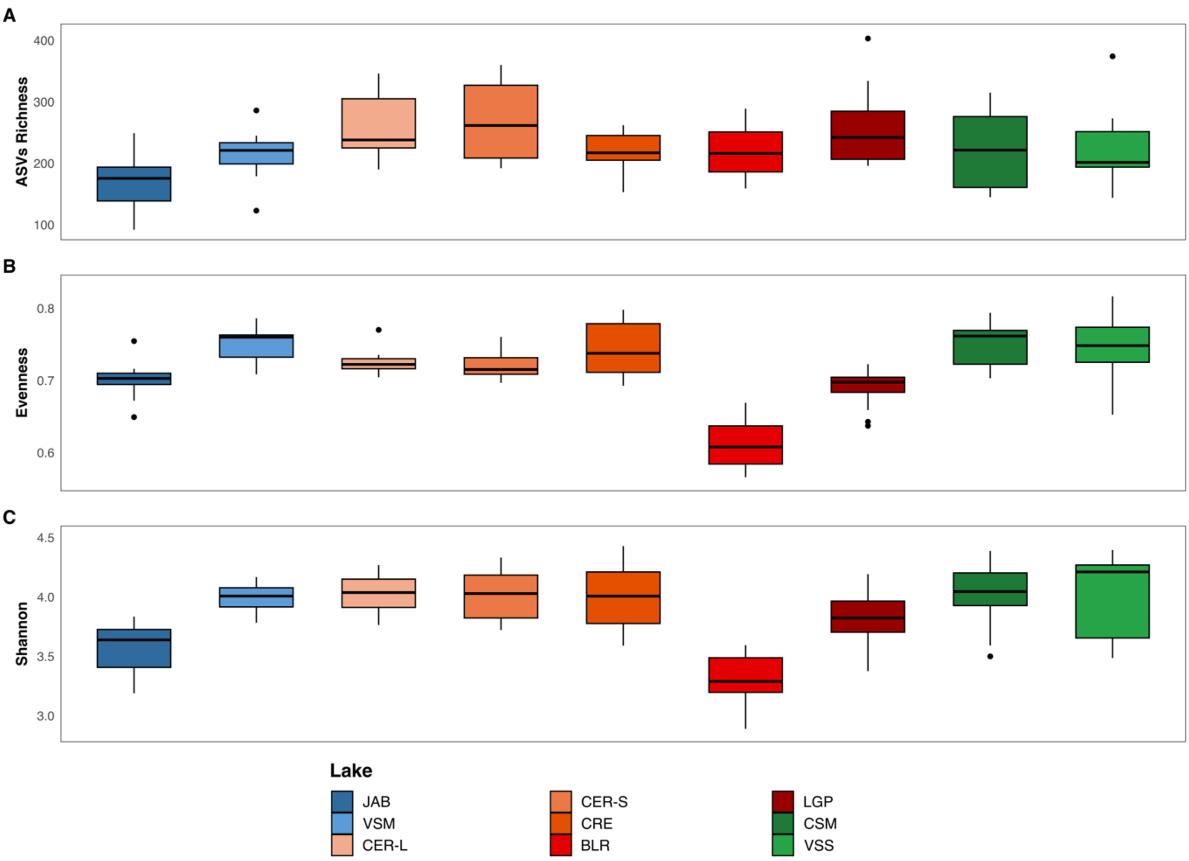


**Fig. S5: Prokaryotic taxa-content alpha-diversity indices.** ASV Richness (**A**), Evenness (**B**) and Shannon diversities (**C**) for each lake over the four summer months (104 samples, **Table S6**). Lakes are colored according to their trophic status (see **Fig. 1B**).

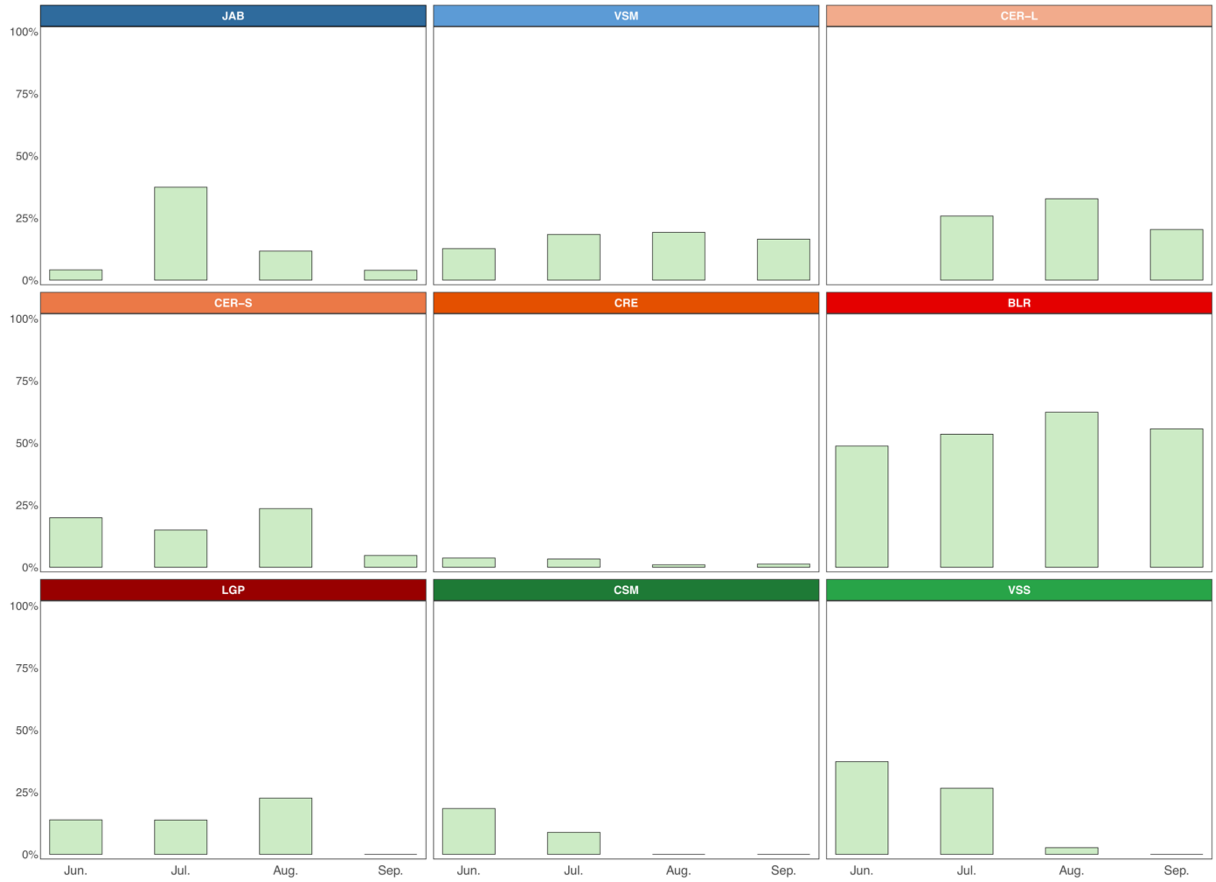


**Fig. S6: Relative abundance (%) of *Cyanobium* ASVs over the four summer months.** Median abundance of the *Cyanobium* ASV (Cyanobacteria), as median percentage of total 16S rRNA reads (104 samples). Lakes are colored according to their trophic status (see **Fig. 1B**).

35

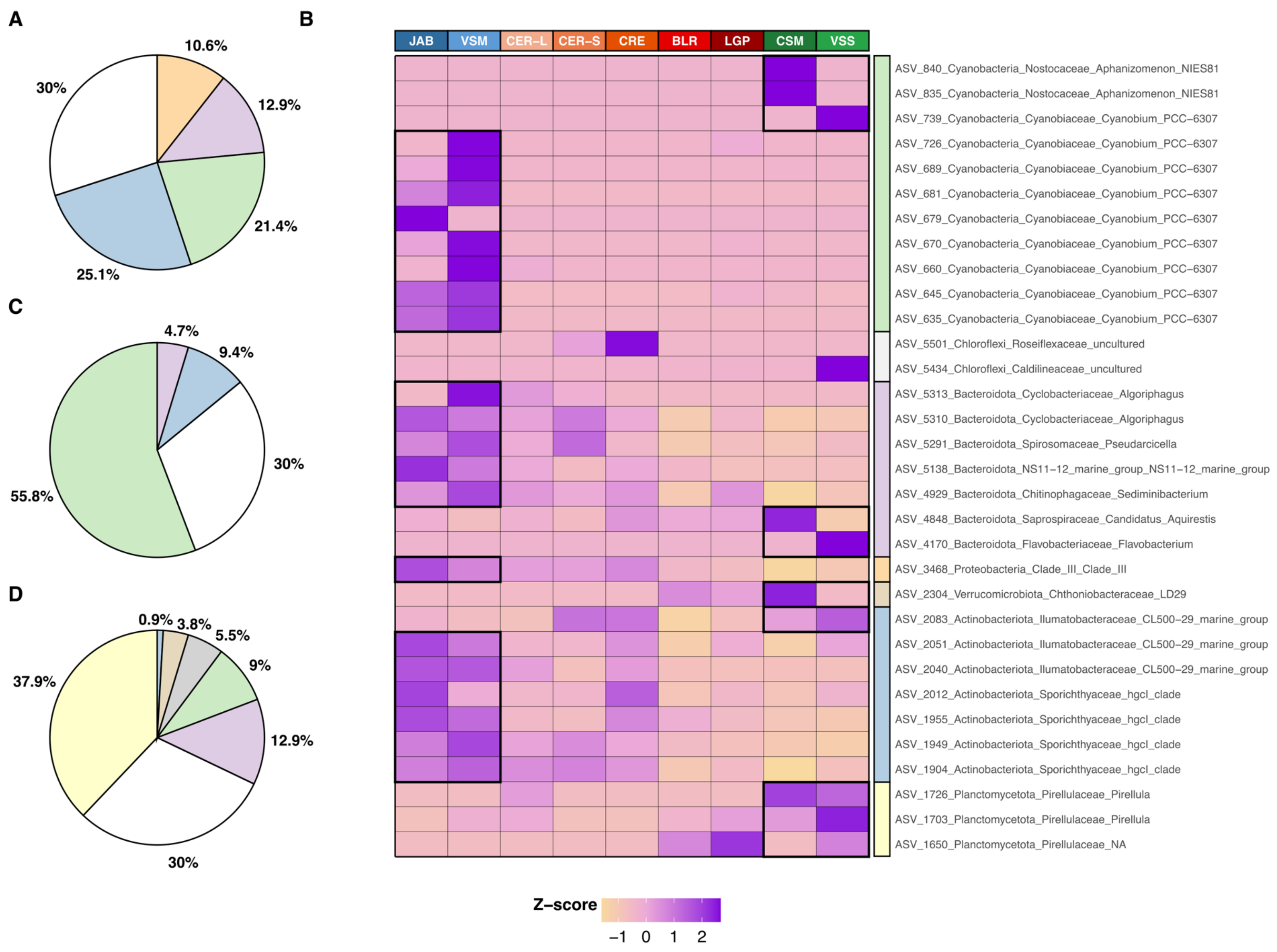

A
10.6%
12.9%
21.4%
25.1%
30%
B
JAB
VSM
CER–L
CER–S
CRE
BLR
LGP
CSM
VSS
ASV_840_Cyanobacteria_Nostocaceae_Aphanizomenon_NIES81
ASV_835_Cyanobacteria_Nostocaceae_Aphanizomenon_NIES81
ASV_739_Cyanobacteria_Cyanobiaceae_Cyanobium_PCC–6307
ASV_726_Cyanobacteria_Cyanobiaceae_Cyanobium_PCC–6307
ASV_689_Cyanobacteria_Cyanobiaceae_Cyanobium_PCC–6307
ASV_681_Cyanobacteria_Cyanobiaceae_Cyanobium_PCC–6307
ASV_679_Cyanobacteria_Cyanobiaceae_Cyanobium_PCC–6307
ASV_670_Cyanobacteria_Cyanobiaceae_Cyanobium_PCC–6307
ASV_660_Cyanobacteria_Cyanobiaceae_Cyanobium_PCC–6307
ASV_645_Cyanobacteria_Cyanobiaceae_Cyanobium_PCC–6307
ASV_635_Cyanobacteria_Cyanobiaceae_Cyanobium_PCC–6307
ASV_5501_Chloroflexi_Roseiflexaceae_uncultured
ASV_5434_Chloroflexi_Caldilineaceae_uncultured
ASV_5313_Bacteroidota_Cyclobacteriaceae_Algoriphagus
ASV_5310_Bacteroidota_Cyclobacteriaceae_Algoriphagus
ASV_5291_Bacteroidota_Spirosomaceae_Pseudarcicella
ASV_5138_Bacteroidota_NS11–12_marine_group_NS11–12_marine_group
ASV_4929_Bacteroidota_Chitinophagaceae_Sediminibacterium
ASV_4848_Bacteroidota_Saprospiraceae_Candidatus_Aquirestis
ASV_4170_Bacteroidota_Flavobacteriaceae_Flavobacterium
ASV_3468_Proteobacteria_Clade_III_Clade_III
ASV_2304_Verrucomicrobiota_Chthoniobacteraceae_LD29
ASV_2083_Actinobacteriota_Ilumatobacteraceae_CL500–29_marine_group
ASV_2051_Actinobacteriota_Ilumatobacteraceae_CL500–29_marine_group
ASV_2040_Actinobacteriota_Ilumatobacteraceae_CL500–29_marine_group
ASV_2012_Actinobacteriota_Sporichthyaceae_hgcI_clade
ASV_1955_Actinobacteriota_Sporichthyaceae_hgcI_clade
ASV_1949_Actinobacteriota_Sporichthyaceae_hgcI_clade
ASV_1904_Actinobacteriota_Sporichthyaceae_hgcI_clade
ASV_1726_Planctomycetota_Pirellulaceae_Pirellula
ASV_1703_Planctomycetota_Pirellulaceae_Pirellula
ASV_1650_Planctomycetota_Pirellulaceae_NA
Z–score
–1 0 1 2
C
4.7%
9.4%
30%
55.8%
D
0.9%
3.8%
5.5%
9%
12.9%
30%
37.9%

**Fig. S7: Set of ASVs that significantly contribute to the ASV-based BC dissimilarities between trophic statuses.** Variance (%) explained by the 34 ASVs with a summed abundance over 0.1% (148 ASVs). AVS are represented at the phylum taxonomic rank, **(A)** mesotrophic *vs.* hypereutrophic; **(C)** eutrophic *vs.* mesotrophic; and **(D)** eutrophic *vs.* hypereutrophic comparison. Slices, colored accordingly to the phylum affiliation (see **Fig. 2B)** represent ASVs that accounted for 70% of the cumulated variance while the other 30% are colored in grey (see Material and Methods**)**. **B**: Relative abundance (Z-score) of the 34 ASVs (of the 0.1% of total reads) significantly explaining the BC dissimilarities between at least one pairwise trophic status comparison. Lakes are colored according to their trophic status (see **Fig. 1B**).

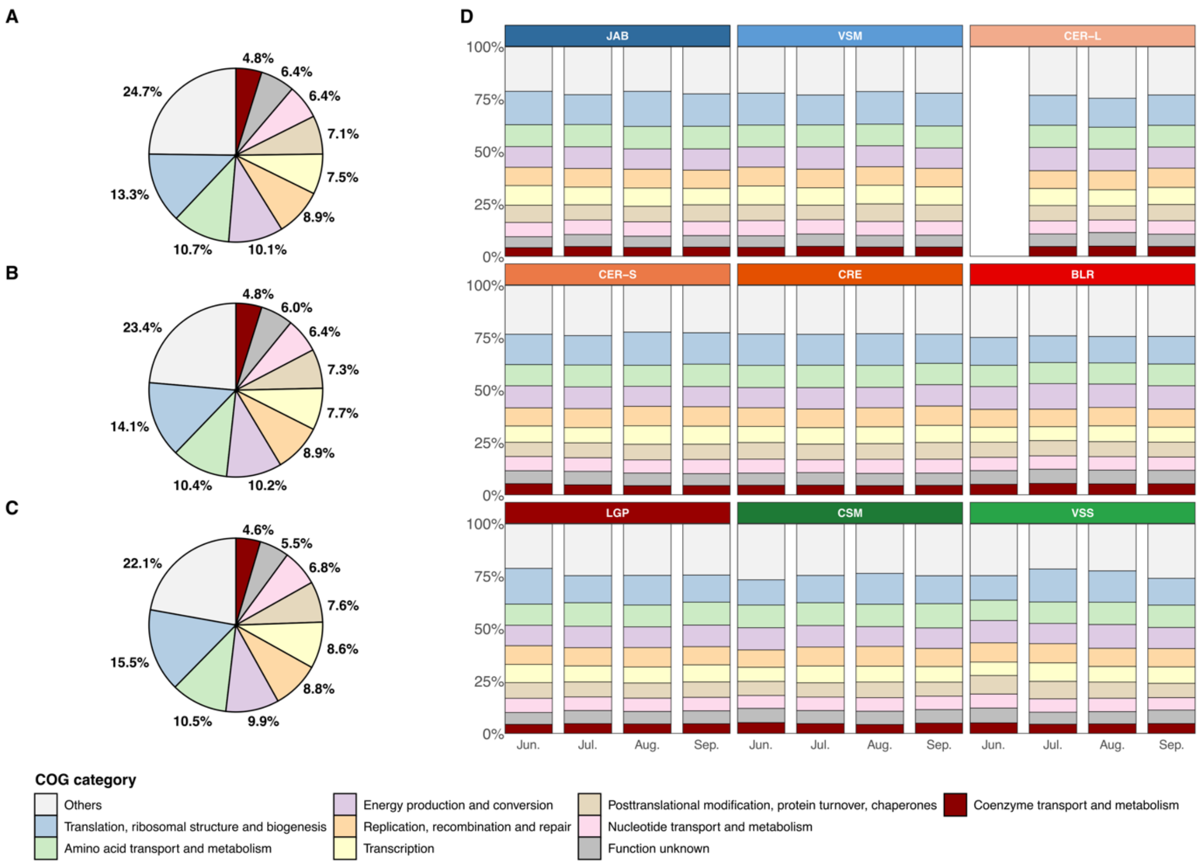


**Fig. S8: Prokaryotic gene-content composition.** COG categories displayed for each trophic status (**A**: hypereutrophic; **B**: eutrophic; **C**: mesotrophic), and by lake and months (**D**), as median proportion of the total KO abundance (n=1 per lake for each month, 35 samples). Only the ten most abundant COG categories are displayed (out of 25, see **Table S9**). Lakes are colored according to their trophic status (see **Fig. 1B**).

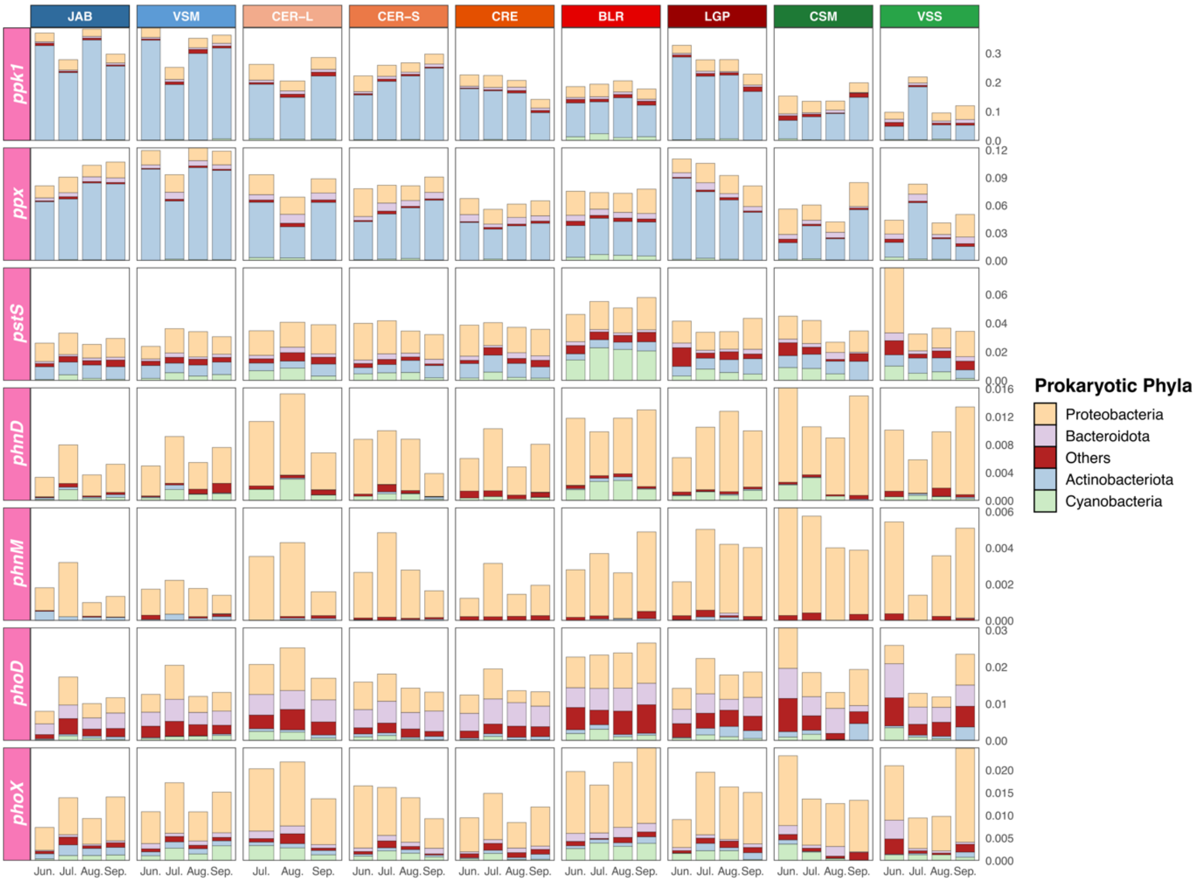


**Fig. S9: Abundance and taxonomic affiliation of the BGC marker genes involved in phosphorus metabolism;** over the four months (as percentage of the total KO abundance, from 0 to 100%; **Table S3**) for each lake (35 samples). Lakes are colored according to their trophic status (see **Fig. 1B**).

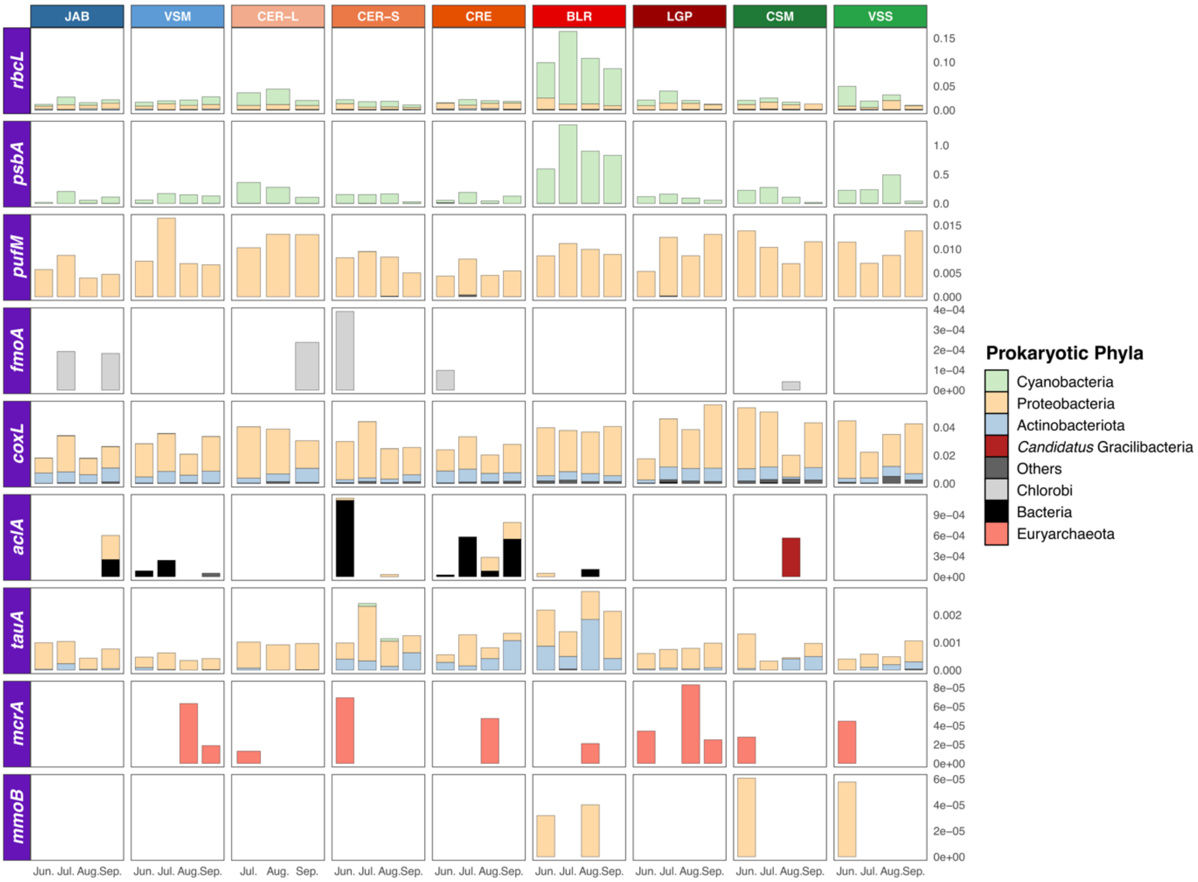


**Fig. S10: Abundance and taxonomic affiliation of the BGC marker genes involved in carbon metabolism;** over the four months (as percentage of the total KO abundance, from 0 to 100%; **Table S3**) for each lake (35 samples). Lakes are colored according to their trophic status (see **Fig. 1B**).

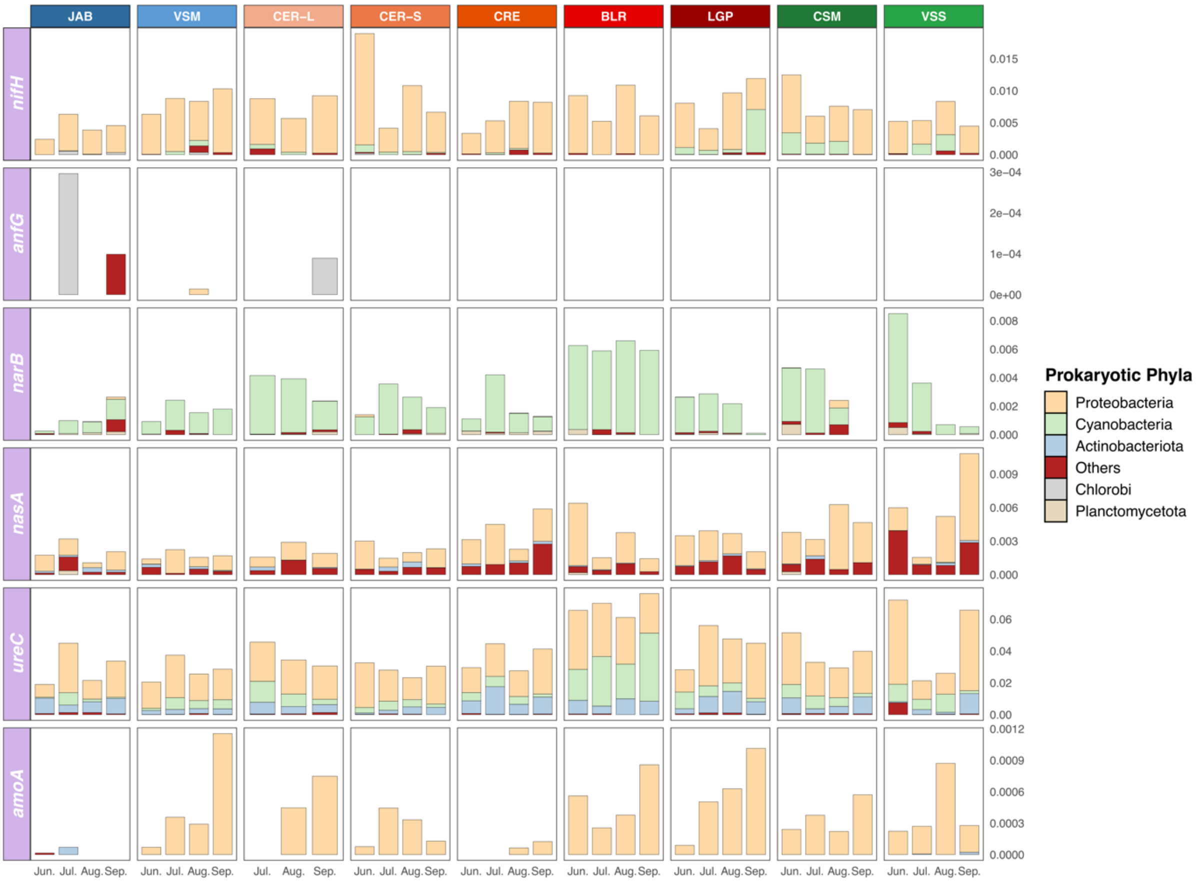


**Fig. S11: Abundance and taxonomic affiliation of the BGC marker genes involved in nitrogen metabolism;** over the four months (as percentage of the total KO abundance, from 0 to 100%; **Table S3**) for each lake (35 samples). Lakes are colored according to their trophic status (see **Fig. 1B**).
